# Brain embeddings with shared geometry to artificial contextual embeddings, as a code for representing language in the human brain

**DOI:** 10.1101/2022.03.01.482586

**Authors:** Ariel Goldstein, Avigail Dabush, Bobbi Aubrey, Mariano Schain, Samuel A. Nastase, Zaid Zada, Eric Ham, Zhuoqiao Hong, Amir Feder, Harshvardhan Gazula, Eliav Buchnik, Werner Doyle, Sasha Devore, Patricia Dugan, Daniel Friedman, Michael Brenner, Avinatan Hassidim, Orrin Devinsky, Adeen Flinker, Uri Hasson

## Abstract

Contextual embeddings, derived from deep language models (DLMs), provide a continuous vectorial representation of language. This embedding space differs fundamentally from the symbolic representations posited by traditional psycholinguistics. Do language areas in the human brain, similar to DLMs, rely on a continuous embedding space to represent language? To test this hypothesis, we densely recorded the neural activity in the Inferior Frontal Gyrus (IFG, also known as Broca’s area) of three participants using dense intracranial arrays while they listened to a 30-minute podcast. From these fine-grained spatiotemporal neural recordings, we derived for each patient a continuous vectorial representation for each word (i.e., a brain embedding). Using stringent, zero-shot mapping, we demonstrated that brain embeddings in the IFG and the DLM contextual embedding space have strikingly similar geometry. This shared geometry allows us to precisely triangulate the position of unseen words in both the brain embedding space (zero-shot encoding) and the DLM contextual embedding space (zero-shot decoding). The continuous brain embedding space provides an alternative computational framework for how natural language is represented in cortical language areas.

## Introduction

Traditionally, investigations into the neural basis of language were based on the symbolic models put forward in psycholinguistics (*1*). These models specify linguistic elements (e.g., nouns, verbs, adjectives, adverbs, etc.) and rule-based operations embedded in hierarchical tree structures. In sharp contrast, *deep language models (DLMs)* trained on massive corpora of natural text provide a radically different family of models for how language is represented in the brain. DLMs encode words as continuous numerical vectors (i.e., contextual embeddings). The geometry of the contextual embedding space (i.e., the relationships between vectors) encodes subtle context-dependent grammatical, semantic, and pragmatic relationships among words (*2–4*). These contextual word embeddings are learned from real-world textual examples “in the wild” without explicit prior knowledge about the structure of language. The fact that continuous contextual embedding spaces can generate well-formed linguistic outputs via vector computations challenges the necessity of discrete, symbolic representations postulated by linguistic (*5*), philosophical (*6*), and psychological (*7*) theories of language.

Recent studies have used contextual word embeddings, derived from DLMs, to successfully model the neural activity measured using fMRI, EEG, MEG, and ECoG during natural speech processing (*8–12*). These studies provide strong evidence that vector-based language representations are useful tools for extracting linguistic information from neural responses. However, it remains unclear whether the neural code within language areas, like DLMs, relies on a vectorial embedding space to represent words during natural language processing. This study provides evidence that the inferior frontal gyrus (IFG, also known as Broca’s area) relies on vector-based brain embeddings to represent words in natural contexts. In particular, the brain embeddings in IFG have strikingly similar geometry to the contextual embeddings learned by DLMs. This result provides a novel, vector-based, computational framework to represent and process language in the human brain.

## Results

Using a dense array of electrocorticographic (ECoG) micro-and macro-electrodes we recorded the neural activity in the IFG (Broca’s area) of three epileptic participants while they listened to a 30-min audio podcast (see Materials and Methods). We focus on IFG as it has been positioned as a central hub for language processing, with an emphasis on semantic and syntactic processing (*1, 9, 13–17*). Overall, we had a dense sampling of 81 electrodes in IFG, with 41, 14, 26 electrodes in participants 1–3 respectively (Fig. 1A). The invasive ECoG recordings provide a dense measure of the neural activity patterns for each word in the IFG. These activity patterns were used to estimate the local brain embedding for each word, where the activity of each electrode serves as a feature (i.e., dimension) in the brain embedding space (Fig. 1D, left). For example, across all participants, we extracted an 81-dimensional brain embedding vector for each word in the story. The brain embeddings were sampled across all electrodes in IFG, which were identified anatomically during surgery, without imposing any additional selection criteria. To assess the selectivity of the results, we also sampled activity from two anatomically adjacent brain regions containing a similar density of electrodes, but not thought to be directly involved in language comprehension: the precentral gyrus and the postcentral gyrus.

**Figure 1.**
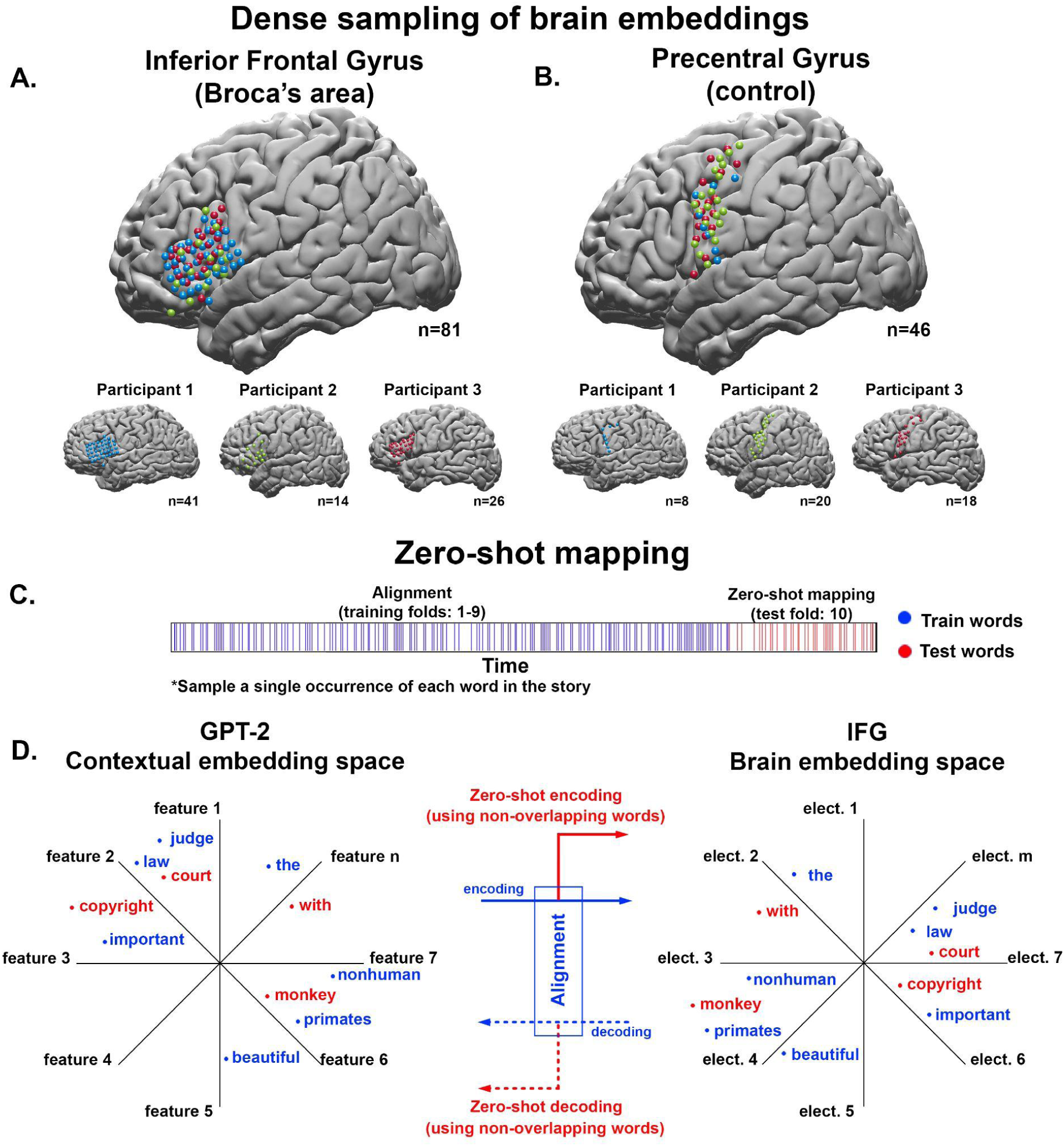
Zero-shot encoding and decoding analysis. (**A**) Dense coverage of the inferior frontal gyrus (Broca’s area) in three patients using micro- and macro-electrodes. Using the Desikan atlas (*18*) we identified electrodes in the left IFG and precentral gyrus in each participant. (**B**) The dense sampling of activity in the adjacent precentral gyrus is used as a control area in the three patients. (**C**) We sampled one instance for each unique word in the podcast. Overall, this resulted in 1100 unique words, which we split into ten folds. Nine folds were used for training (blue) and one fold, containing 110 unique, non-overlapping words, was used for testing (red). This procedure was repeated across all folds. (**D**, left) We extracted the contextual embeddings from GPT-2 for each of the words. Using PCA, we reduced the contextual embeddings to 50-dimensional vectors (50 features). (**D**, right) We used the dense sampling of activity patterns across electrodes in IFG to estimate a brain embedding vector for each of the 1100 words. The brain embeddings were extracted within each participant and across participants. (**D**, center) We used nine of the folds to align the brain embeddings (multi-electrode activity patterns) with the GPT-2 contextual embeddings for each word in the training set. The solid blue arrow denotes the alignment phase of the encoding analysis, in which we align the contextual embeddings to the brain embeddings based on a subset of training words; the solid red arrow denotes the evaluation phase of the encoding analysis, where we predict brain embeddings for novel words from the contextual embeddings. The dotted blue arrow denotes the alignment phase of the decoding analysis, in which we align the brain embeddings to the contextual embeddings based on a subset of training words; the dotted red arrow denotes the evaluation phase of the decoding analysis, where we predict contextual embeddings for novel words from the brain embeddings.

We used zero-shot mapping, a stringent analysis method in machine learning, to demonstrate that IFG brain embeddings have shared geometrical properties with contextual embeddings derived from a high-performing DLM (GPT-2). Zero-shot mapping tests the ability of the model to predict (or interpolate) the brain embeddings (encoding) or contextual embeddings (decoding) of words the model was not exposed to during training. The zero-shot analysis imposes a stringent separation between the words used for aligning the brain embeddings and contextual embeddings and the words used for evaluating the mapping (red and blue words in Fig. 1D). We sampled one instance of each unique word in the podcast, resulting in 1100 words with no repetitions. Each of those 1100 unique words is represented by a 1600-dimensional contextual embedding extracted from the final layer of GPT-2. The contextual embeddings were reduced to 50-dimensional vectors using PCA (Materials and Methods). We then divided these 1100 words into 10 contiguous folds, with 110 unique words in each segment. Crucially, there was no overlap between the words in each fold. We used nine of the folds to align the brain embeddings derived from IFG (Broca’s area) with the 50-dimension contextual embeddings derived from GPT-2 (Fig. 1D, blue words). The alignment between the contextual embeddings and brain embeddings was done separately for each lag (at 200 ms resolution; see Materials and Methods) within an 8-second window (4 s before and 4 s after the onset of each word, where lag 0 is word onset). The remaining words in the non-overlapping test fold were used to evaluate the zero-shot mapping (Fig 1D, red words).

### Zero-shot encoding

In zero-shot encoding analysis, we used nine folds of the data (990 unique words) to learn a linear transformation between the contextual embeddings from GPT-2 and the brain embeddings from IFG. Next, we used the tenth fold to predict (interpolate) IFG brain embeddings for a new set of 110 words to which the encoding model was never exposed. Critically, relying on a single instance of each word for predicting an unseen word forces the zero-shot mapping to rely solely on the information embedded in the geometry of the contextual embedding space. For example, we used the words “important”, “law”, “judge”, “nonhuman”, etc, to align the contextual embedding space to the brain embedding space. Using the alignment model (encoding model), we next predicted the brain embeddings for a new set of words “copyright”, “court” and “monkey”, etc. Accurately predicting IFG brain embeddings for the unseen words is viable only if the geometry of the brain embedding space matches the geometry of the contextual embedding space. If the geometric relationships among words are not shared between the brain embeddings and contextual embeddings, learning the mapping on one set of words cannot accurately predict the neural activity for a new, non-overlapping set of words.

In the zero-shot encoding analysis, we successfully predicted brain embeddings in IFG for words that were not seen during training (Fig. 2A, blue lines). We correlated each of the predicted brain embeddings with the actual brain embedding in the test fold and averaged the correlations across words in the test fold (separately for each lag). The averaged correlations of the unseen words were significant at multiple time points surrounding word onset for all three participants (for details on the significance test, see Materials and Methods), with peak correlations roughly 200 ms after word onset. Furthermore, the encoding performance for unseen words was significant up to −700 ms before word onset, which provides evidence for the engagement of IFG in context-based next-word prediction (*9*). The zero-shot mapping results were robust in each individual participant and the group level (Fig. 2B, blue lines).

**Figure 2.**
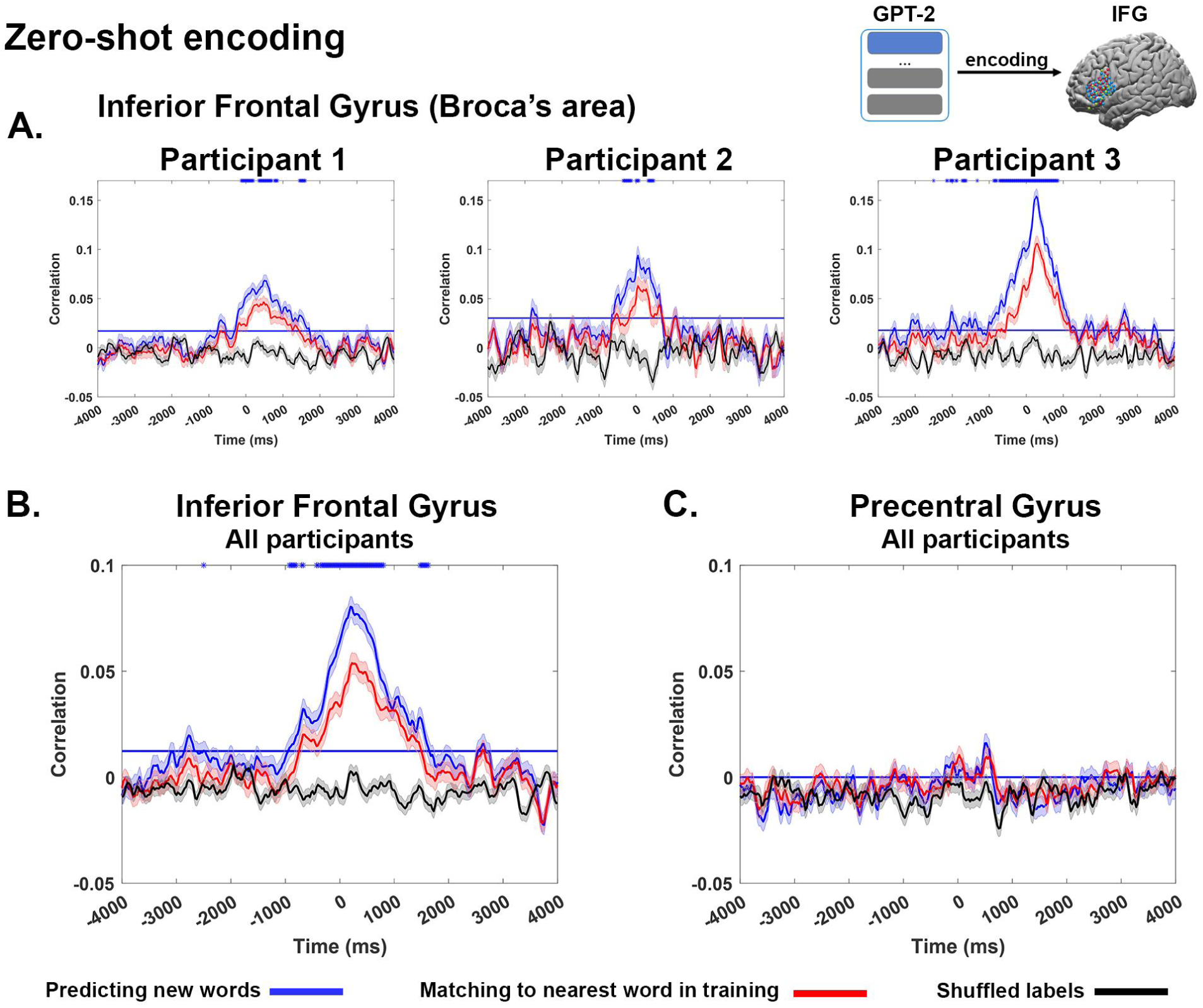
Encoding analysis reveals shared geometry between contextual embeddings and brain embeddings. (**A**) Zero-shot encoding between the contextual embeddings and the brain embeddings in IFG for each individual patient. The solid blue line shows the average correlation between the predicted brain embeddings and actual brain embeddings in IFG for all words across all test sets. Significant correlations peak after word onset, but also precede word onset (the significance threshold marked by the horizontal blue line). The red line shows the zero-shot encoding for the word from the training set that is most similar (nearest neighbor) to each test word. Note, that the reduced correlations for the nearest training embeddings indicate that the zero-shot mapping is able to accurately interpolate to new embeddings not seen during the training phase (at the level of individual patients). The blue asterisks represent a significant difference (FDR corrected) between the correlation with the actual contextual embeddings (blue line) and the correlation with the nearest embedding from the training set (red line). The black line shows the zero-shot encoding between shuffled contextual embeddings and brain embeddings. (**B**) Zero-shot encoding for brain embeddings was extracted across all participants. (**C**) Zero-shot encoding for electrodes sampled from the anatomically adjacent control area, precentral gyrus. No alignment between brain embeddings and contextual embeddings was observed in this non-linguistic area.

The zero-shot encoding performance was close to zero when we randomly matched the words in the test-fold with mismatching contextual embeddings (Fig. 2AB, black lines). Furthermore, the predicted neural activity pattern (brain embedding) for each of the unseen words was highly selective to the IFG and was completely absent in an adjacent control area, the precentral gyrus (Fig. 2C, blue lines). To ensure that the lack of zero-shot mapping in the precentral gyrus is not due to the lower spatial sampling, we replicated the findings in individual participants, which have a comparable number of electrodes in the precentral gyrus and IFG (Fig. S1). We also combined electrodes in the precentral gyrus and postcentral gyrus to ensure that the increase in sampling density does not improve the zero-shot mapping in control areas (Fig. S2).

### Precise neural interpolation based on shared geometry

The zero-shot encoding analysis demonstrates that the shared geometry between contextual embeddings and brain embeddings in IFG is sufficient to predict the neural activation patterns for unseen words. A possible confound, however, is the intrinsic co-similarities among word representations in both spaces. For example, the embedding for the word “monkey” may be similar to the embedding for the word “baboon” (in most contexts); it is also likely that the activation patterns for these words in the IFG are similar (*19, 20*). To control for this, we devised a control analysis to determine whether the zero-shot mapping learns to precisely interpolate the brain embedding for unseen words using the shared geometry across both embedding spaces. We repeated the zero-shot mapping analysis—however, instead of using the actual contextual embedding for each of the unseen words in the test fold, we used the contextual embedding of the closest word in the training folds (based on cosine similarity). For example, instead of using the contextual embedding for “baboon” (a previously-unseen word in the test fold), we used the contextual embedding of the most similar word in the training set (“monkey”). If the zero-shot analysis simply matches the predicted brain embedding with the nearest similar contextual embedding from the language model among the training words, the switch to the nearest training embedding will not deteriorate the results. In contrast, if the alignment truly exposes shared geometry between the two embedding spaces, using the embedding for the nearest training word will significantly reduce the zero-shot encoding performance. Indeed, the results showed a significant decrease in performance when using the nearest training embeddings (Fig. 2, red line). This is evident both at the group level and in each individual participant. This further supports the claim that alignment between the contextual and brain embeddings revealed fine-grained shared geometric structure between the two representational spaces.

### Zero-shot decoding of individual words

In the zero-shot encoding analysis, we successfully mapped the contextual embeddings into the brain embedding space of the IFG. Next, we reversed the mapping (i.e., decoding analysis) to map the brain embedding space into the contextual embedding space of GPT-2 (Fig. 1D). For this purpose, we used a two-step classification procedure (*9*). First, we trained a convolutional neural network to align the brain embedding of each word in the training folds to their corresponding contextual embedding (see Materials and Method and Appendix I). Next, we used the trained neural network to predict the contextual embeddings for the signal associated with each unseen word in the test fold. The cosine distance between the predicted contextual embedding and the actual contextual embedding was used to classify the nearest word in the 110-word test set (i.e., zero-shot decoding). As with the encoding analysis, we repeated this procedure separately for different temporal shifts relative to work onset. We used the area under the receiver operating characteristic curve (ROC-AUC) to quantify the amount of information for each word. ROC-AUC of 0.5 indicates chance performance, and ROC-AUC of 1 indicates perfect classification among all test words.

Using zero-shot decoding, we were able to classify words well above chance level (Fig. 3). We were able to decode words at the group level and also to replicate the results in all three individuals. Peak classification was observed at a lag of roughly 320 ms after onset with a ROC-AUC of 0.60, 0.65, and 0.67 in individual participants and 0.70 at the group level (Fig. 3, pink line). Shuffling the labels flattened the ROC-AUC to 0.5 (chance level, Fig. 3 black lines). Running the same procedure on the precentral gyrus control area (Fig. 3, green line) yielded AUC closer to chance level (maximum AUC of 0.55). The residual classification in the precentral gyrus may be attributed to the proximity to the IFG, or to the enhanced power of deep nonlinear models for learning the alignment between embedding spaces.

**Figure 3.**
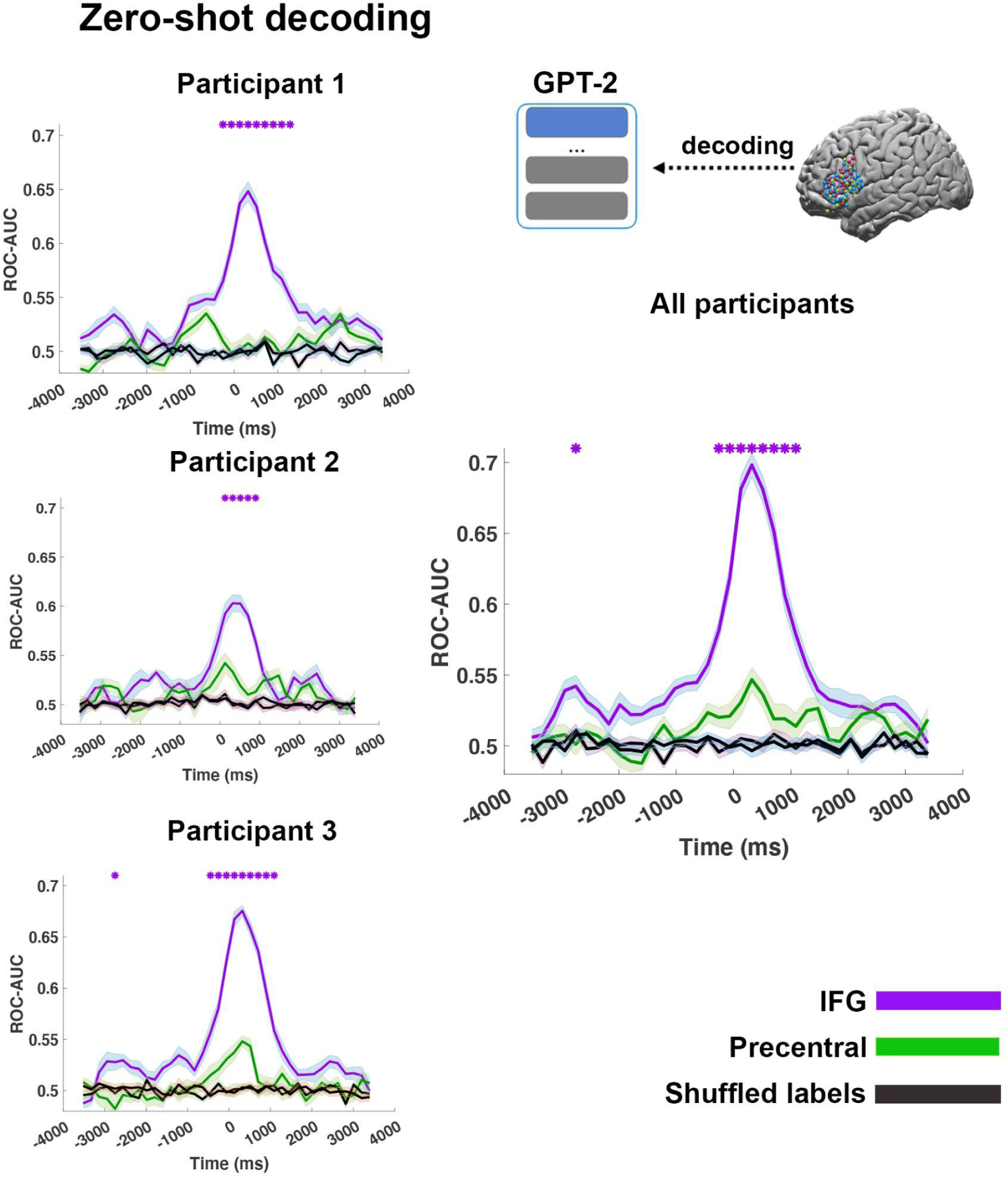
Zero-shot decoding of unseen words after aligning contextual embeddings with the brain embeddings of inferior frontal gyrus (IFG) and precentral gyrus. Average ROC-AUCs zero-shot word classification performance across folds based on the predicted brain embeddings in IFG (pink line) and precentral gyrus (green line). Zero-shot decoding was performed in each individual participant and using brain embeddings extracted across all participants. The classification is performed for all unseen words in each test fold and performance is averaged across all 10 test folds (the error-bars are across the folds). The graph shows the average zero-shot classification performance based on the cosine distance of each predicted embedding and all other 110 words in the test fold. The performance of word classification is measured by the area under the receiver operating characteristic curve (ROC-AUC). Zero-shot classification peaks after word onset, but may precede word onset by up to 1000 ms.The black lines show the zero-shot classification between brain embeddings and shuffled contextual embeddings. In pink asterisks we mark the significant difference between IFG embedding and precentral embedding, using paired sample t-test of the AUCs and bonferroni correction for multiple comparisons..

## Discussion

How does the brain encode the compositional and contextual meaning of words in natural language? Our findings sharply depart from the symbolic representations and rule-based syntactic operations of classical neurolinguistics. Using dense, spatiotemporal ECoG recordings, we sampled the continuous vector representation (brain embeddings) of words in a natural narrative within a well-localized language area, the IFG (also known as Broca’s area). Using zero-shot encoding and decoding, we demonstrate that, similar to DLMs, IFG relies on continuous vectorial embeddings to represent words in natural contexts. We were able to encode and decode at the group level, as well as at the individual level, which attests to the robustness of this neural representational format. Importantly, the brain embedding space shares a strikingly similar geometry with the contextual embedding space learned by DLMs. This shared geometry was sufficient to predict the activity patterns in IFG to a new set of words not seen during training. This effect diminished in an adjacent, non-linguistic control area. Critically, the zero-shot predictions were precise enough to predict (interpolate) the location of novel words rather than to regress to the semantically nearest word from the training set.

While we discovered a shared geometry between brain embeddings derived from IFG and contextual embedding derived from DLM, our analyses do not assess the dimensionality of both embedding spaces (*21*). In this work, we reduce the dimensionality of the contextual embeddings from 1600 to 50-dimensions. In this lower dimension, we demonstrate a shared continuous-vectorial geometry between both embedding spaces. To assess the dimensionality of the brain embeddings representation in IFG, we need a denser sampling of both the underlying neural activity and the semantic space of natural language. Similarly, to assess the dimensionality of the contextual embeddings space, additional analyses of the feature dependencies are needed (*21*).

A fundamental question in neurolinguistics is whether language processing relies on symbolic representation and rule-based computations or whether language processing relies on continuous vectorial space. For most classical psycholinguistics, the prior encoding and decoding results (*19, 20, 22, 23*) are impressive methodological achievements in reading semantic information from neural activity (i.e., “mind-reading”). However, they and other decoding results (*24, 25*) do not support the strong claim that the brain relies on continuous vector space to code and process language. In contrast, our dense sampling of the brain embedding space in IFG, a well-localized language area, provides direct evidence that brain embeddings are viable computational means to code and process each word as it is embedded in real-life natural narratives. Thus, our results provide a new computational framework for how language is coded and processed in the human brain (*2, 3, 26*).

Why is the embedded representational space for coding language shared between DLMs and the human brain? After all, there are fundamental differences between the way DLMs and the human brain learn a language. For example, DLMs are trained on massive text corpora, whereas children learn spoken language from interacting with their local community of native speakers. Furthermore, current DLMs rely on the transformer architecture, which is not biologically plausible (*27*). Although DLMs rely on very different mechanisms for learning, we recently found evidence for three shared computational principles between autoregressive DLMs and the human brain (*9*). Importantly, deep language models should be viewed as statistical learning models which learn the structure of language by conditioning the contextual embeddings on the way humans use words in natural contexts. If humans, like DLMs, learn the structure of language from the way language is used by speakers, then the two representational spaces should converge (*21, 28*). Indeed, recent work has begun to show how implicit knowledge about syntactic and compositional properties of language are embedded in the contextual representations of deep language models (*29, 30*). The shared representational space suggests that the human brain, like DLMs, relies on overparameterized optimization to learn the statistical structure of language from other speakers in the natural world (*28*).

While the geometry of brain’s and DLMs embeddings is similar, it is certainly not identical. Future work is needed to more thoroughly explore the differences. For example, the linguistic inputs to which humans are exposed may have a different statistical structure from the linguistic inputs used to train DLMs, resulting in different embedding spaces. While most DLMs (such as GPT2, BERT, T5) are trained on large corpora of text, children learn a language in the context of open-ended multimodal verbal interaction with their social environment. This suggests that DLMs trained on audiovisual conversations may learn contextual embeddings that better fit the brain embeddings in the human brain. In addition, differences between the contextual and brain embedding spaces may arise from the differences in the circuit architecture, learning rule, and objective function, all of which are only loosely related to the human brain (*28, 31*).

To conclude, the remarkable alignment between brain embeddings and DLM contextual embeddings, combined with accumulated evidence across recent papers (*8–11, 21*) suggests that the brain may rely on vector-space representation and computation to capture the richness of natural language. The shift from a symbolic representation of language to a vectorial representation derived from a statistical learning model is a paradigm shift for understanding the neural basis of language processing in the human brain.

## Materials and Methods

### Participants

Three patients (2 female; 24–48 years old) with treatment-resistant epilepsy undergoing intracranial monitoring with subdural grid and strip electrodes for clinical purposes participated in the study. Three of the study participants consented to have an FDA-approved hybrid clinical-research grid implanted that includes additional electrodes in between the standard clinical contacts. The hybrid grid provides a higher spatial coverage without changing clinical acquisition or grid placement. Each participant provided informed consent in accordance with protocols approved by the New York University Grossman School of Medicine Institutional Review Board. Patients were informed that participation in the study was unrelated to their clinical care and that they could withdraw from the study at any point without affecting their medical treatment.

### Stimulus

Participants were presented with a 30-minute auditory story stimulus, “So a Monkey and a Horse Walk Into a Bar: Act One, Monkey in the Middle” taken from the *This American Life* podcast. The onset of each word was marked using the Penn Phonetics Lab Forced Aligner (*32*) and manually validated and adjusted (if necessary). The stimulus and alignment process are described in prior work (*9*).

### Data acquisition and preprocessing

A total of 1106 electrodes were placed on the left hemispheres and 233 on the right hemispheres (signal sampled at 512 Hz). The full description of the ECoG recording procedure is provided in prior work (*9*). Electrode-wise preprocessing consisted of four main stages: First, large spikes exceeding 4 quartiles above and below the median were removed and replacement samples were imputed using cubic interpolation. Second, the data were re-referenced using common average referencing. Third, 6-cycle wavelet decomposition was used to compute the high-frequency broadband (HFBB) power in the 70–200 Hz band, excluding 60, 120, 180 Hz line noise. In addition, the HFBB time series of each electrode was log-transformed and z-scored. Fourth, the signal was smoothed using a Hamming window with a kernel size of 50 ms. The filter was applied in both the forward and reverse directions in order to maintain the temporal structure. Additional preprocessing details can be found in prior work (*9*).

### Contextual embedding

We extracted contextualized word embeddings from GPT-2 using the Hugging Face environment (*33*). We first converted the words from the raw transcript (including punctuation and capitalization) to tokens comprising either whole words or sub-words (e.g., there’s → there ‘s). We used a sliding window of 1024 tokens, moving one token at a time, to extract the embedding for the final word in the sequence (i.e., the word and its history). We extracted the activity of the final hidden layer of GPT-2 (which has 48 hidden layers). The contextual embedding of a word is the activity of the last hidden layer given all the words up to and not including the word of interest (note, that in GPT-2 the word is predicted using the last hidden state).

### Brain embedding

We extracted brain embedding for specific ROIs by averaging the neural activity in a 200 ms window for each electrode in the ROI. This means that if there are *N* electrodes in a specific ROI for a specific patient, then, for each lag around onset (ranging from −4 sec to +4 sec in 25 ms shifts), there will be an *N*-dimensional embedding, where each feature is the averaged neural activity of a specific electrode of the neural recordings in a window of 200 ms (102 time points) centered at the lag.

### Zero-shot encoding model

Linear encoding models were estimated at each lag relative to word onset in order to predict the brain embedding for each word from the corresponding contextual embedding. Prior to fitting the encoding model, we smoothed the signal using a rolling 200-ms window. We used a 10-fold cross-validation procedure ensuring that for each cross-validation fold, the model was estimated from a subset of unique training words and evaluated on a nonoverlapping subset of unique, held-out test words: the words and their corresponding brain embeddings were split into a training set (90% of the unique words) for model estimation and a test set (10% of the unique words) for model evaluation (zero-shot analysis). For each cross-validation fold, we used ordinary least squares (OLS) multiple linear regression to estimate a weight vector (for the 50-dimensional model feature space) based on the training words. We then used those weights to predict the neural responses at each electrode (comprising a “brain embedding” across electrodes) for the test words. We evaluated model performance by computing the correlation between the predicted brain embedding and the actual brain embedding (i.e., the distributed neural activity pattern) for each held-out test word; we then averaged these correlations across test words. This procedure was repeated in full at 321 lags at 25-ms increments from −4000 ms to 4000 ms relative to word onset. As a control analysis, for each test word, we instead used the nearest contextual embedding (in terms of cosine distance) from the training set to predict the brain embedding for the test word.

### Statistical significance

We used a bootstrap hypothesis test to assess the statistical significance of the correlations between the predicted brain embeddings and actual brain embeddings. The test statistic reported for each lag is the average of the correlations between the predicted brain embedding and actual brain embedding across all test words. We then resampled these correlations across words with replacement (5000 bootstrap samples) to generate a bootstrap distribution around the mean correlation. We then computed a p-value based on the null hypothesis that the correlation is zero. This procedure was repeated for each lag (321 lags) and we controlled the false discovery rate (FDR) at *q* = .01 (*34*).

In order to test whether there was a significant difference between the performance of the model using the actual contextual embedding for the test words compared to the performance using the nearest word from the training fold, we performed a permutation test. At each iteration, we permuted the differences in performance across words and assigned the mean difference to a null distribution. We then computed a p-value for the difference between the test embedding and the nearest training embedding based on this null distribution. This procedure was repeated to produce a p-value for each lag and we corrected for multiple tests using FDR.

In order to compare the difference between classifier performance using IFG embedding or precentral embedding for each lag we used paired sample t-test. For each lag we compared the AUC of each word classified with the IFG embedding or precentral embedding. Thank we used bonferroni correction to account for the multiple comparison.

### Zero-shot decoding model

We used a decoding model to classify unseen words from the corresponding brain embeddings. The neural signals were first averaged per electrode in 10 62.5-ms bins spanning 625 ms for each lag. Each bin consisted of 32 data points (the neural recording sampling rate was 512 Hz). We used a 10-fold cross-validation procedure ensuring that for each cross-validation fold, the decoding model was trained on a subset of unique training words and evaluated on a nonoverlapping subset of unique, held-out test words: the words and their corresponding brain embeddings were split into a training set (90% of the unique words) for model estimation and a test set (10% of the unique words) for model evaluation (zero-shot analysis). A neural network decoder (see architecture in Appendix I) was trained to predict the contextual embedding for each word from the corresponding brain embedding at a specific lag. Eight training folds were used for training the decoder (training set), one fold was used for early stopping (development set), and one fold was used to assess model generalization (test set). The neural net was optimized to minimize the mean squared error (MSE) when predicting the embedding.

The classification was performed in two ways: First, we computed the cosine similarity between the predicted contextual embedding and all the unique contextual embeddings in the dataset (Fig. 3 blue lines). We used a softmax transformation on these scores (logits). For each label, we used these logits to evaluate whether the decoder predicted the matching word and computed an ROC-AUC for the label. In this evaluation strategy, each test word is evaluated against the other test words in that particular test set. To improve the performance of the decoder, we implemented an ensemble of models. We independently trained 6 classifiers with randomized weight initializations and randomized the batch order supplied to the neural net for each lag. This procedure generated 6 predicted embeddings. Thus, for each predicted embedding, we repeated the distance calculation from each word label 6 times. These 6 values were averaged and used to compute the ROC-AUC.

### Code availability

The code will be available upon acceptance of publication

---

**Figure S1.**
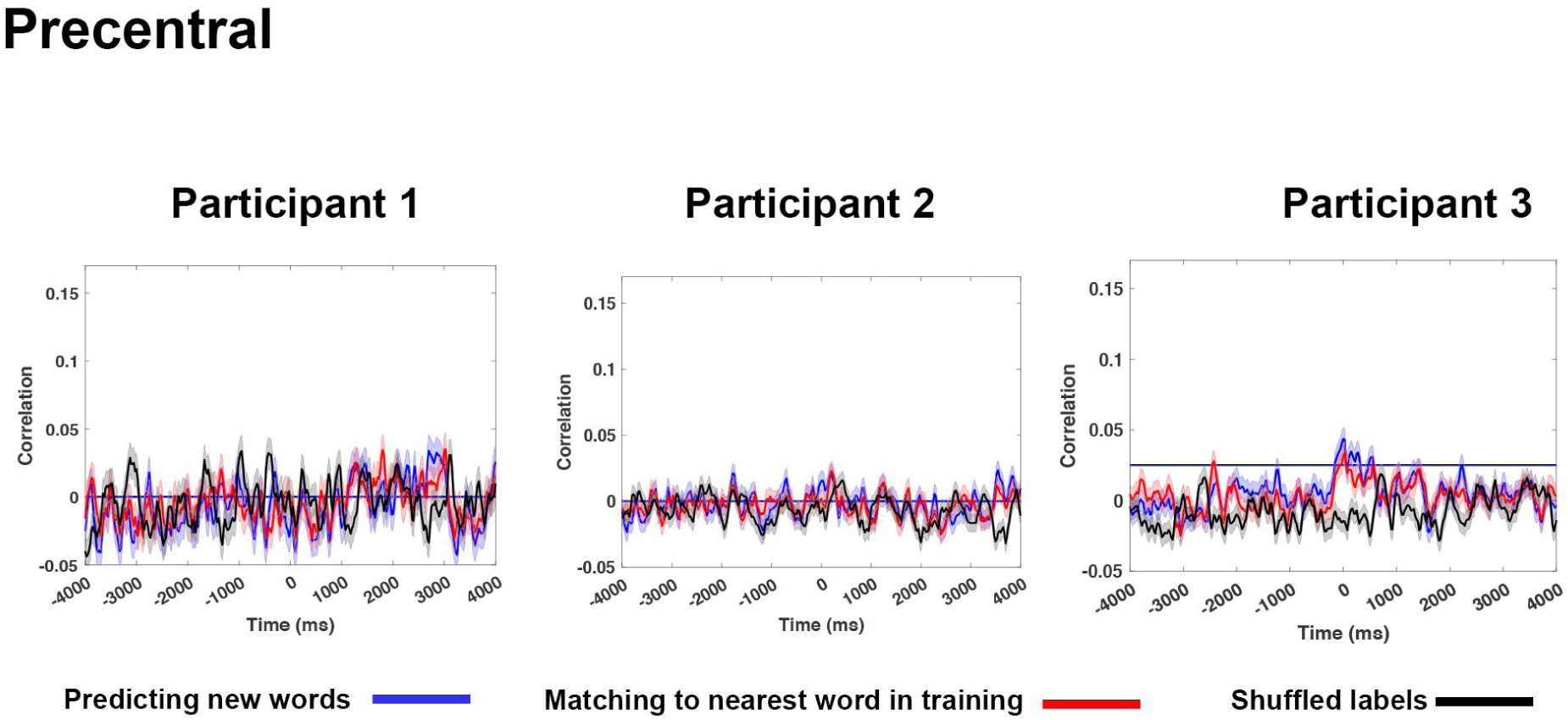
Zero-shot encoding between the contextual embeddings and the brain embeddings of the precentral gyrus (control area) for each patient. Zero-shot encoding is evaluated at each lag (−4000 to 4000 ms). None of the encoding models found a significant difference between near neighbor and the actual word.

**Figure S2.**
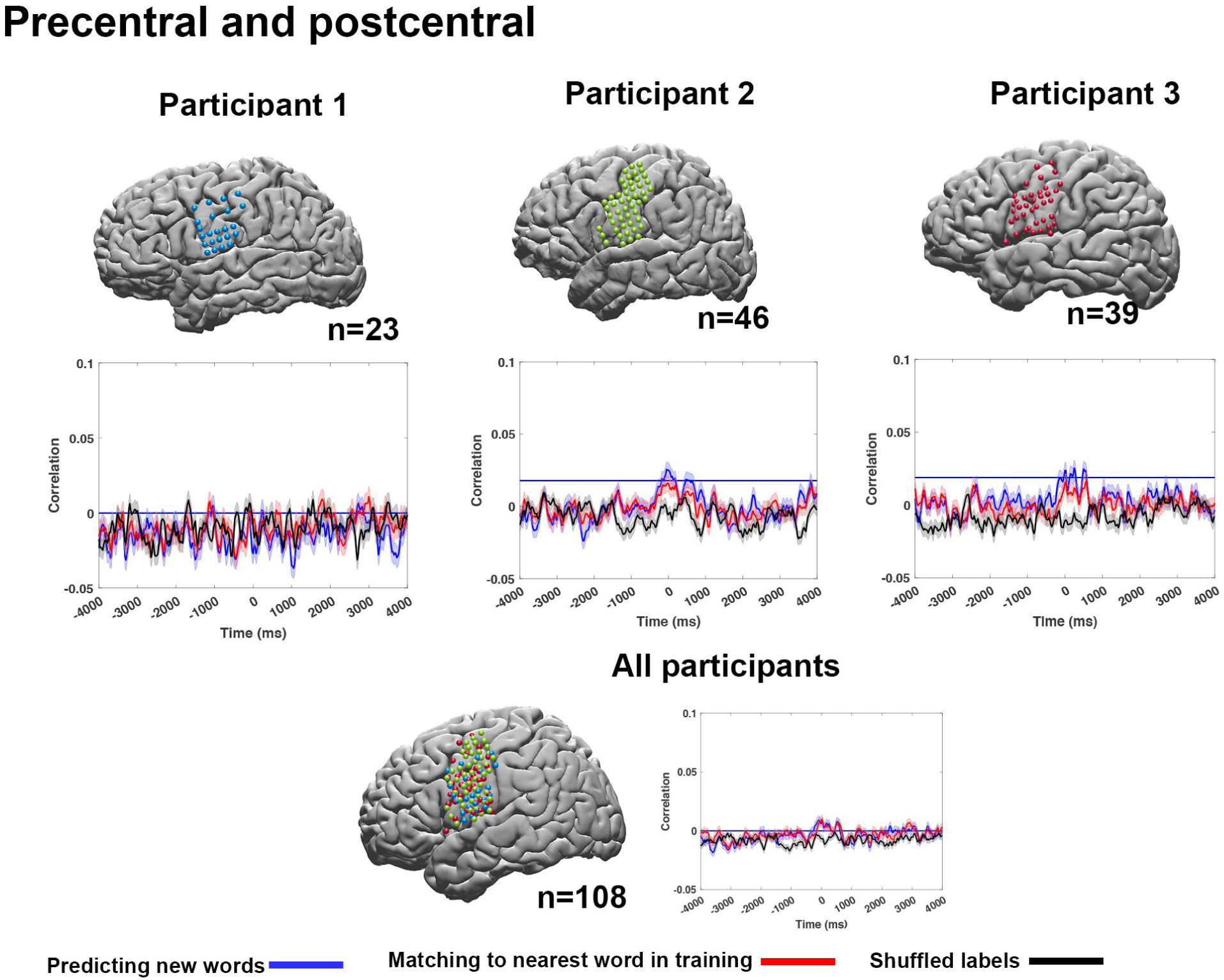
In order to control for the increased number of electrodes in the IFG we collapsed the precentral and postcentral gyrus and re-run the encoding analysis. Implementing the same encoding procedures and significant testing did not yield any lag that showed significant difference between the original test-set and the near neighbor.

## Appendix I Decoding architecture

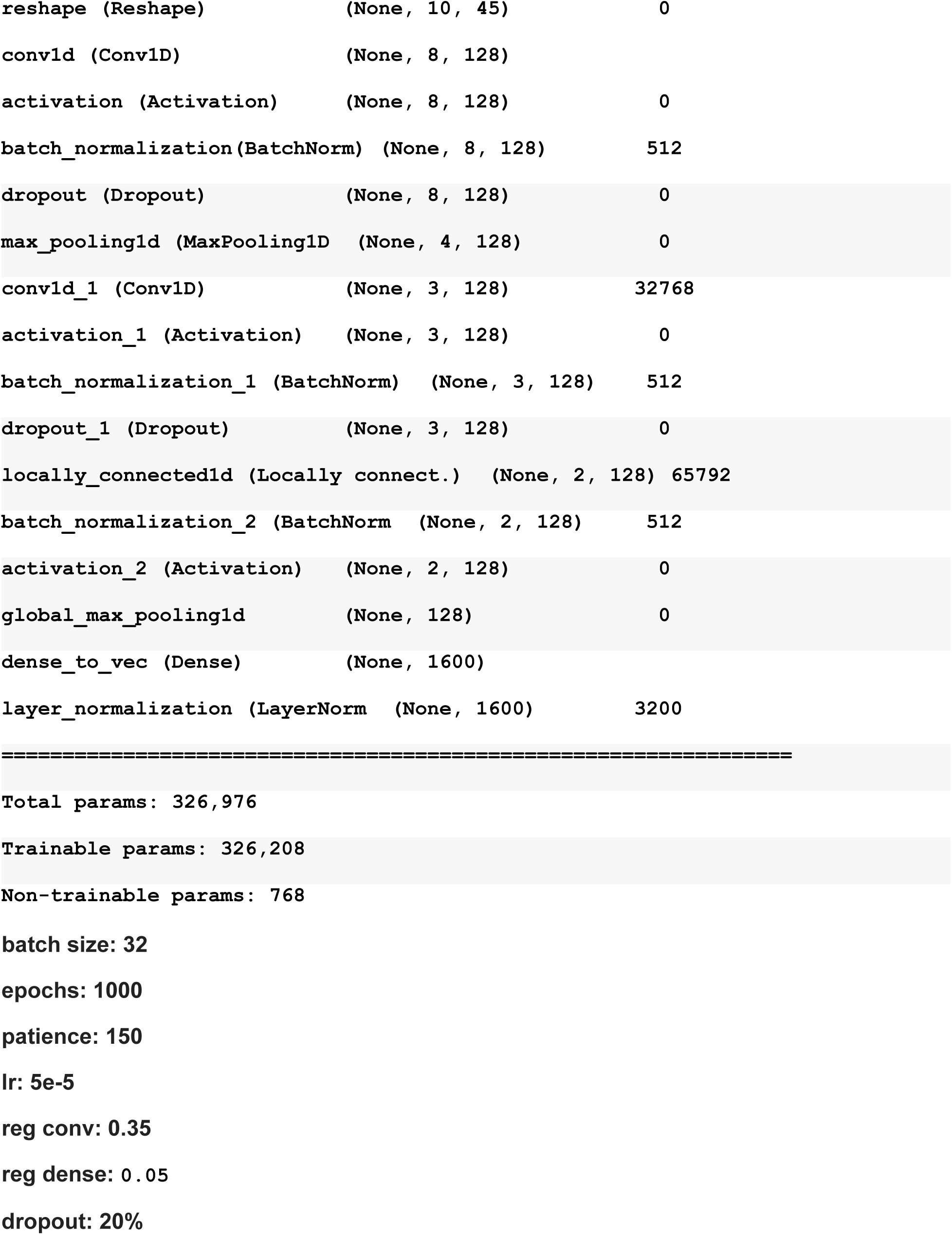

## References

1. P. Hagoort, On Broca, brain, and binding: a new framework. Trends Cogn. Sci. 9, 416–423 (2005).

2. A. Radford, J. Wu, R. Child, D. Luan, D. Amodei, I. Sutskever, Language models are unsupervised multitask learners. OpenAI blog. 1, 9 (2019).

3. T. B. Brown, B. Mann, N. Ryder, M. Subbiah, J. Kaplan, P. Dhariwal, A. Neelakantan, P. Shyam, G. Sastry, A. Askell, S. Agarwal, A. Herbert-Voss, G. Krueger, T. Henighan, R. Child, A. Ramesh, D. M. Ziegler, J. Wu, C. Winter, C. Hesse, M. Chen, E. Sigler, M. Litwin, S. Gray, B. Chess, J. Clark, C. Berner, S. McCandlish, A. Radford, I. Sutskever, D. Amodei, Language Models are Few-Shot Learners. arXiv [cs.CL] (2020), (available at http://arxiv.org/abs/2005.14165).

4. Z. Yang, Z. Dai, Y. Yang, J. Carbonell, R. R. Salakhutdinov, Q. V. Le, in Advances in Neural Information Processing Systems, H. Wallach, H. Larochelle, A. Beygelzimer, F. d\textquotesingle Alché-Buc, E. Fox, R. Garnett, Eds. (Curran Associates, Inc., 2019; https://proceedings.neurips.cc/paper/2019/file/dc6a7e655d7e5840e66733e9ee67cc69-Paper.pdf), vol. 32.

5. N. Chomsky, Approaching UG from Below. Interfaces Recursion = Language?, pp. 1–30.

6. J. A. Fodor, The Language of Thought (Harvard University Press, 1975).

7. J. L. McClelland, T. T. Rogers, The parallel distributed processing approach to semantic cognition. Nat. Rev. Neurosci. 4, 310–322 (2003).

8. M. Schrimpf, I. A. Blank, G. Tuckute, C. Kauf, E. A. Hosseini, N. Kanwisher, J. B. Tenenbaum, E. Fedorenko, The neural architecture of language: Integrative modeling converges on predictive processing. Proc. Natl. Acad. Sci. U. S. A. 118 (2021), doi:10.1073/pnas.2105646118.

9. A. Goldstein, Z. Zada, E. Buchnik, M. Schain, A. Price, B. Aubrey, S. A. Nastase, A. Feder, D. Emanuel, A. Cohen, A. Jansen, H. Gazula, G. Choe, A. Rao, C. Kim, C. Casto, L. Fanda, W. Doyle, D. Friedman, P. Dugan, R. Reichart, S. Devore, A. Flinker, L. Hasenfratz, A. Hassidim, M. Brenner, Y. Matias, K. A. Norman, O. Devinsky, U. Hasson, Thinking ahead: spontaneous prediction in context as a keystone of language in humans and machines,, doi:10.1101/2020.12.02.403477.

10. C. Caucheteux, A. Gramfort, J. R. King, GPT-2’s activations predict the degree of semantic comprehension in the human brain. bioRxiv (2021) (available at https://www.biorxiv.org/content/10.1101/2021.04.20.440622v2.abstract).

11. M. Toneva, L. Wehbe, in 33rd Conference on Neural Information Processing Systems (NeurIPS 2019), Vancouver, Canada. (2019; http://arxiv.org/abs/1905.11833).

12. C. Caucheteux, J.-R. King, Brains and algorithms partially converge in natural language processing. Commun Biol. 5, 134 (2022).

13. P. Hagoort, P. Indefrey, The neurobiology of language beyond single words. Annu. Rev. Neurosci. 37, 347–362 (2014).

14. X. Yang, H. Li, N. Lin, X. Zhang, Y. Wang, Y. Zhang, Q. Zhang, X. Zuo, Y. Yang, Uncovering cortical activations of discourse comprehension and their overlaps with common large-scale neural networks. NeuroImage. 203 (2019), p. 116200.

15. B. Ishkhanyan, V. Michel Lange, K. Boye, J. Mogensen, A. Karabanov, G. Hartwigsen, H. R. Siebner, Anterior and Posterior Left Inferior Frontal Gyrus Contribute to the Implementation of Grammatical Determiners During Language Production. Front. Psychol. 11, 685 (2020).

16. L. L. LaPointe, Paul Broca and the Origins of Language in the Brain (Plural Publishing, 2012).

17. D. Saur, B. W. Kreher, S. Schnell, D. Kümmerer, P. Kellmeyer, M.-S. Vry, R. Umarova, M. Musso, V. Glauche, S. Abel, W. Huber, M. Rijntjes, J. Hennig, C. Weiller, Ventral and dorsal pathways for language. Proc. Natl. Acad. Sci. U. S. A. 105, 18035–18040 (2008).

18. R. S. Desikan, F. Ségonne, B. Fischl, B. T. Quinn, B. C. Dickerson, D. Blacker, R. L. Buckner, A. M. Dale, R. P. Maguire, B. T. Hyman, M. S. Albert, R. J. Killiany, An automated labeling system for subdividing the human cerebral cortex on MRI scans into gyral based regions of interest. Neuroimage. 31, 968–980 (2006).

19. T. M. Mitchell, S. V. Shinkareva, A. Carlson, K.-M. Chang, V. L. Malave, R. A. Mason, M. A. Just, Predicting human brain activity associated with the meanings of nouns. Science. 320, 1191–1195 (2008).

20. F. Pereira, B. Lou, B. Pritchett, S. Ritter, S. J. Gershman, N. Kanwisher, M. Botvinick, E. Fedorenko, Toward a universal decoder of linguistic meaning from brain activation. Nat. Commun. 9, 963 (2018).

21. R. Antonello, J. Turek, V. Vo, A. Huth, Low-Dimensional Structure in the Space of Language Representations is Reflected in Brain Responses. arXiv [cs.CL] (2021), (available at http://arxiv.org/abs/2106.05426).

22. A. G. Huth, W. A. de Heer, T. L. Griffiths, F. E. Theunissen, J. L. Gallant, Natural speech reveals the semantic maps that tile human cerebral cortex. Nature. 532, 453–458 (2016).

23. J. G. Makin, D. A. Moses, E. F. Chang, Machine translation of cortical activity to text with an encoder-decoder framework. Nat. Neurosci. 23, 575–582 (2020).

24. J. V. Haxby, A. C. Connolly, J. S. Guntupalli, Decoding neural representational spaces using multivariate pattern analysis. Annu. Rev. Neurosci. 37, 435–456 (2014).

25. N. Kriegeskorte, P. K. Douglas, Interpreting encoding and decoding models. Curr. Opin. Neurobiol. 55, 167–179 (2019).

26. C. Raffel, N. Shazeer, A. Roberts, K. Lee, S. Narang, M. Matena, Y. Zhou, W. Li, P. J. Liu, Exploring the limits of transfer learning with a unified text-to-text transformer. arXiv preprint arXiv:1910.10683 (2019) (available at https://www.jmlr.org/papers/volume21/20-074/20-074.pdf).

27. D. J. Heeger, K. O. Zemlianova, A recurrent circuit implements normalization, simulating the dynamics of V1 activity. Proc. Natl. Acad. Sci. U. S. A. 117, 22494–22505 (2020).

28. U. Hasson, S. A. Nastase, A. Goldstein, Direct Fit to Nature: An Evolutionary Perspective on Biological and Artificial Neural Networks. Neuron. 105, 416–434 (2020).

29. J. Hewitt, C. D. Manning, in Proceedings of the 2019 Conference of the North American Chapter of the Association for Computational Linguistics: Human Language Technologies, Volume 1 (Long and Short Papers) (2019), pp. 4129–4138.

30. T. Linzen, M. Baroni, Syntactic Structure from Deep Learning. Annu. Rev. Appl. Linguist. (2021), doi:10.1146/annurev-linguistics-032020-051035.

31. B. A. Richards, T. P. Lillicrap, P. Beaudoin, Y. Bengio, R. Bogacz, A. Christensen, C. Clopath, R. P. Costa, A. de Berker, S. Ganguli, C. J. Gillon, D. Hafner, A. Kepecs, N. Kriegeskorte, P. Latham, G. W. Lindsay, K. D. Miller, R. Naud, C. C. Pack, P. Poirazi, P. Roelfsema, J. Sacramento, A. Saxe, B. Scellier, A. C. Schapiro, W. Senn, G. Wayne, D. Yamins, F. Zenke, J. Zylberberg, D. Therien, K. P. Kording, A deep learning framework for neuroscience. Nat. Neurosci. 22, 1761–1770 (2019).

32. J. Yuan, M. Liberman, Speaker identification on the SCOTUS corpus. J. Acoust. Soc. Am. 123, 3878 (2008).

33. L. Tunstall, L. von Werra, T. Wolf, Natural Language Processing with Transformers: Building Language Applications with Hugging Face (O’Reilly Media, 2022).

34. Y. Benjamini, Y. Hochberg, Controlling the False Discovery Rate: A Practical and Powerful Approach to Multiple Testing. Journal of the Royal Statistical Society: Series B (Methodological). 57 (1995), pp. 289–300.

